# Mapping single-cell mechanics in the early embryogenesis of *Xenopus laevis* using atomic force microscopy

**DOI:** 10.1101/2025.07.11.664270

**Authors:** Miki Yamamoto, Takayoshi Yamamoto, Takahiro Kotani, Yuki Miyata, Tatsuo Michiue, Takaharu Okajima

## Abstract

During early embryonic development, cells undergo rapid cleavage divisions accompanied by morphological changes driven by mechanical cues. However, the spatiotemporal mechanics of regulative embryonic cells remains poorly understood. Here, we use atomic force microscopy (AFM) to map single-cell stiffness of *Xenopus laevis* embryos, a model of regulative development, from early cleavage (stage 6) to the onset of gastrulation (stage 11). To ensure stable AFM mapping, vitelline membrane-removed embryos were immobilized in custom grooved agarose wells and gently held using a dulled glass pipette. AFM observations revealed marked mechanical heterogeneity within the animal hemisphere: stiffness in apical cytoplasmic regions, which are defined as the central apical surface excluding cell-cell boundaries, varied among cells, regardless of size, indicating intrinsic variability. Cell-cell boundaries consistently showed high stiffness, suggesting strong adhesion, as commonly observed in epithelial monolayers in vitro. In contrast, in the vegetal hemisphere during gastrulation, cell-cell boundaries exhibited low stiffness, indicative of weaker adhesion. Additionally, micron-scale stiff inclusions were detected in the apical vegetal cytoplasm, with sizes comparable to those of yolk platelets. These findings demonstrate the capability of AFM to probe the microscale mechanical architecture of developing regulative embryos and to uncover regional mechanical asymmetries between the animal and vegetal hemispheres. Such asymmetries may contribute to key morphogenetic processes during early vertebrate development.

## 1. Introduction

During the early stages of embryogenesis, cells undergo profound morphological and functional changes, leading to the emergence of complex multicellular architecture. These transitions are increasingly recognized to be influenced not only by genetic and biochemical factors but also by mechanical cues (*1-10*). Therefore, direct mechanical measurements of developing embryos have become indispensable for elucidating the physical principles underlying early developmental processes. To date, the mechanical properties of embryos have been investigated at multiple spatial scales (*11-15*). Whole-embryo stiffness has been assessed using compression-based methods, in which embryos are deformed between parallel plates (*16, 17*). Microfluidic probes have enabled localized mechanical measurements within tissues (*15, 18-21*), while microaspiration and laser ablation techniques have provided critical insights into tensile forces at the single-cell level (*12, 22*). Recent technological advances allow for unprecedented spatiotemporal resolution in mapping cellular mechanics during early cleavage stages, particularly in mosaic embryos such as ascidians, through high-precision methodologies including atomic force microscopy (AFM) (*23, 24*) and Brillouin microscopy (*25-27*).

The mechanical architecture of embryonic cells arises from the integrated interplay between the actomyosin-rich cortex and the underlying viscoelastic cytoplasm. The cortex forms a thin, contractile layer beneath the plasma membrane, while the cytoplasm is a gel-like composite of cytosol, organelles, and cytoskeletal elements. These two compartments, forming a continuous dynamic system, can be probed by AFM as they deform under mechanical stress, although their interconnected nature complicates the analysis of their contributions to overall cellular stiffness (*28*).

Among species being regulative embryos, *Xenopus laevis* has been widely used as a model organism in developmental mechanobiology (*11, 29-31*). Previous studies on *Xenopus laevis* embryos demonstrated that AFM enables not only the surface morphology of the embryo (*32*), but also the mechanical properties of the embryos (*33-38*), brain tissue (*31, 35, 39, 40*), and oocytes (*41, 42*). However, how single-cell stiffness in regulative embryos changes during the developmental stages remains to be characterized. Here, using AFM, we investigated the spatiotemporal mechanical properties of *Xenopus laevis* embryos during early embryogenesis from stage 6 (cleavage stage) to stage 11 (gastrula stage) (Fig. 1a). The AFM mapping revealed both intra-and intercellular stiffness in developing *Xenopus laevis* embryos at the single-cell level. A cell-to-cell variability in mechanical properties was observed in the animal hemisphere. Around the gastrula stage, distinct differences in mechanical profile were observed between animal and vegetal regions. Furthermore, AFM could detect the yolk platelets underneath the cell membrane, which were observed as stiffer granules.

**FIGURE 1.**
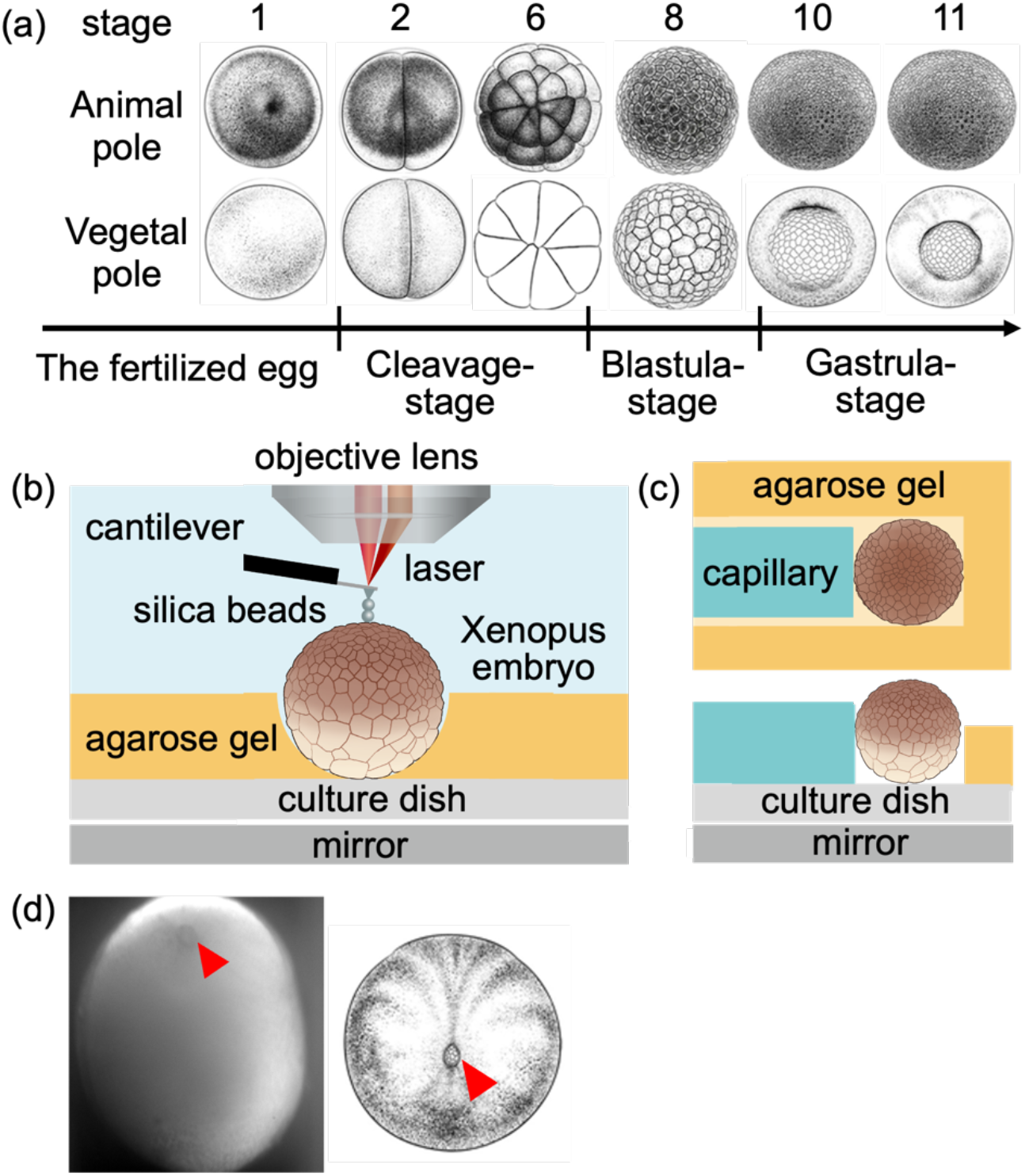
(a) Schematic morphological change in *Xenopus laevis* embryo during the early embryogenesis from the fertilized egg to stage 11 (mid-gastrula) from *Xenopus* illustrations©Natalya Zahn (2022) (*48*). (b) Schematic of AFM with an upright optical microscope. *Xenopus* embryo was immobilized in a semi-cylindrical groove agarose wells and cultured in 1/10× Steinberg’s solution. The underside morphology of the embryo was observed through a mirror placed beneath the gel plate, using the optical microscope. (c) Schematic of embryo placed in a semi-cylindrical well in a top view (upper figure) and side view (lower figure). The embryo was slightly contacted on the edge of the agarose well (orange) and then gently held using a dulled glass capillary (green). (d) A microscope image of a developing embryo at around stage 12, viewed from the vegetal pole using the mirror. The yolk plug of the embryo is denoted by the arrowhead. After checking the stage with the image, the animal hemisphere was measured by AFM.

## 2. Experimental methods

### 2.1 Manipulation of *Xenopus laevis* embryos

Unfertilized eggs were obtained from female *Xenopus laevis* injected with gonadotropin and artificially fertilized with testis homogenate. Fertilized eggs were de-jellied with 4.6% L-cysteine-HCl solution (pH 7.8) and incubated in 1/10× Steinberg’s solution at about 17 ºC. The vitelline membrane of embryos was manually removed with tweezers (*37*).

A semi-cylindrical-shaped agarose well was prepared by inserting a glass capillary (1.2 mm in outer diameter) into the pre-gel solution before gelation in a culture dish. After the agarose had solidified, the capillary was carefully removed from the gel to form the well. Vitelline membrane-removed embryos were subsequently positioned within semi-cylindrical agarose wells and gently secured between the agarose wall and a dulled glass capillary (Fig. 1c). This configuration was essential for stable mapping of the vegetal hemisphere, which shows lower buoyancy than the animal hemisphere.

### 2.2 AFM measurements and analysis

A custom-built atomic force microscope was employed to simultaneously map the relative surface height *H* and apparent Young’s modulus *E* of *Xenopus* embryos (Fig. 1b). The atomic force microscope system was integrated with an upright optical microscope (Eclipse FN1; Nikon) with a liquid-immersion objective (plan fluor, ×10, Nikon), enabling optical lever detection, as reported previously (*23, 24, 43-45*). Force measurements were performed using a rectangular cantilever (Biolever-mini, BL-AC40TS-C2; Olympus) with a nominal spring constant below 0.1 N/m. Before the experiments, the spring constant of the cantilever was calibrated in aqueous conditions via the thermal noise method (*46, 47*). To ensure a well-defined probe-sample contact geometry and eliminate undesired interaction between the cantilever and the sample surface, more than two silica microspheres (radius *R* of ca. 3.5 μm) were fixed in tandem to the cantilever tip using epoxy resin (Fig. 1b) (*23, 24, 45*). The AFM force mapping experiments were conducted on a specific area of the surface of *Xenopus* embryos at about 23°C.

A piezoelectric scanner (P-563.3CD; Physik Instruments), operated via LabVIEW-FPGA software (National Instruments), was used for stage positioning. The scan area was manually adjusted across developmental stages to accommodate morphological changes, with scan widths within 200 μm and lateral step sizes of 3 μm, depending on sample geometry. During AFM measurements targeting the animal hemispheres during the gastrula stage, the morphological features of the vegetal hemisphere were visualized via a mirror mounted beneath the sample dish (Fig. 1d). This setup facilitated the precise staging of the embryos based on morphological criteria, especially in the vegetal region (*48*).

The *E* of embryonic cells was estimated from force-distance curves obtained during the approach phase using a modified Hertz contact model (*49*). The AFM probe consisted of a spherical indenter (radius *R*) contacting the sample surface at a local tilt angle *θ* and indentation depth *δ*, yielding a loading force *F* governed by the relation:

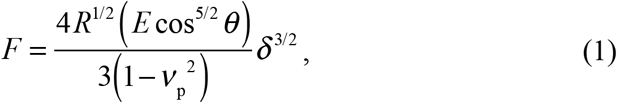

where *ν*_p_ is the Poisson’s ratio of the cell, assumed to be 0.5, and *θ* of the sample surface was estimated from AFM topographic (relative height *H*) maps. The modified Hertz contact model was valid for *θ* < 40° in the case that *E* was < ∼10 kPa (*49*). Thus, we discussed the phenomena observed in mapping regions that satisfied *θ* < 40°. To facilitate visualization, the *E* maps were log-scaled using log_10_ (*E* [Pa]), whereas the *H* maps were presented on a linear scale. For all analyses, the maximum indentation depth was restricted to be less than 2.5 μm, depending on developmental stages.

## 3. Results

### 3.1 Stiffness in the animal hemisphere at the early cleavage stage

In *Xenopus laevis*, cleavage divisions proceed rapidly after fertilization but exhibit differences in cell size between the animal and vegetal regions from stage 4. Even at stage 6, cells in the vegetal hemisphere, which reserve substantial yolks that impede cytokinesis, retain a large, rounded morphology. This geometry restricts access by the AFM probe to the apical surface of vegetal hemisphere cells. Therefore, we initially focused on the animal hemisphere at stage 6, where probe contact enabled mechanical mapping of the developing embryo (Fig. 2a). Since embryo samples were de-jellied and stripped of the vitelline membrane before mechanical mapping, the whole embryo morphology reproducibly exhibited a flattened elliptical geometry, deviating from their native spherical form. Despite their deformations, we confirmed that most embryos continued their normal developmental progression even after AFM experiments.

**FIGURE 2.**
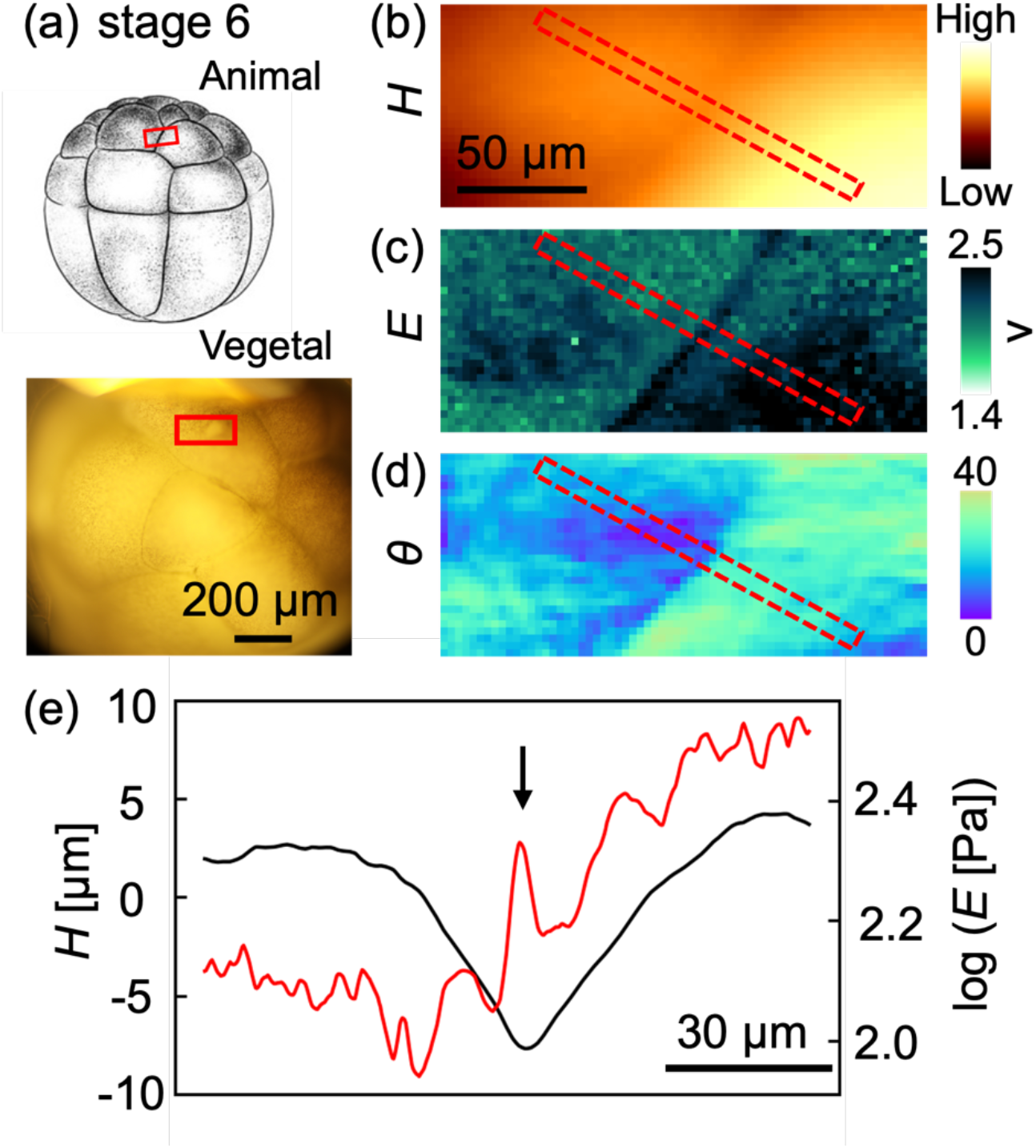
(a) Schematic morphology (upper) and an optical microscopic image (lower) of a *Xenopus laevis* embryo at stage 6. AFM-measured region is indicated by an open rectangle (red). The AFM mapping images of *H* (b) and *E* (c), and *θ* (d) estimated from (b). (e) Plots of averaged *H* (c) and *E* (d) values in the region denoted by open dashed-line rectangles (b-d, red). The topography is plotted by flattening the tilting of the height image. The arrow represents the position of the cell-cell boundary.

The AFM-based spatial mapping of relative height, *H* (Fig. 2b) and apparent Young’s modulus, *E* (Fig. 2c) revealed that while the height maps appeared relatively smooth across the apical surface (Fig. 2b, e), the stiffness landscape exhibited marked heterogeneity (Fig. 2c, e), suggesting that localized variations in intracellular mechanical properties result from structural differences beneath the cell membrane, rather than from surface topography of the membrane. The stiffness was remarkably high in the cell-cell boundaries (Fig. 2c, e), suggesting tight cell-cell interactions. Importantly, the cell stiffness in certain regions in the apical cytoplasmic region, which is the area near the center of the apical cell surface excluding the cell-cell boundaries, was higher than that at the cell-cell boundaries (Fig. 2e). These findings highlight the presence of intrinsic intracellular mechanical complexity in embryonic cells at the early cleavage stage.

### 3.2 Stiffness in the animal hemisphere at the midblastula stage

We next performed mechanical mapping of *Xenopus laevis* embryos at stage 8 (midblastula stage) (Fig. 3a) at three time points (see legend of Fig. 3 for details). At this stage, we again observed that cells exhibited marked stiffening along cell-cell boundaries (Fig. 3b-d), as observed at stage 6. We also observed that cell stiffness within apical cytoplasmic regions displayed considerable temporal variations between neighboring cells (Fig. 3e), reflecting the intrinsic mechanical heterogeneity of cells.

**FIGURE 3.**
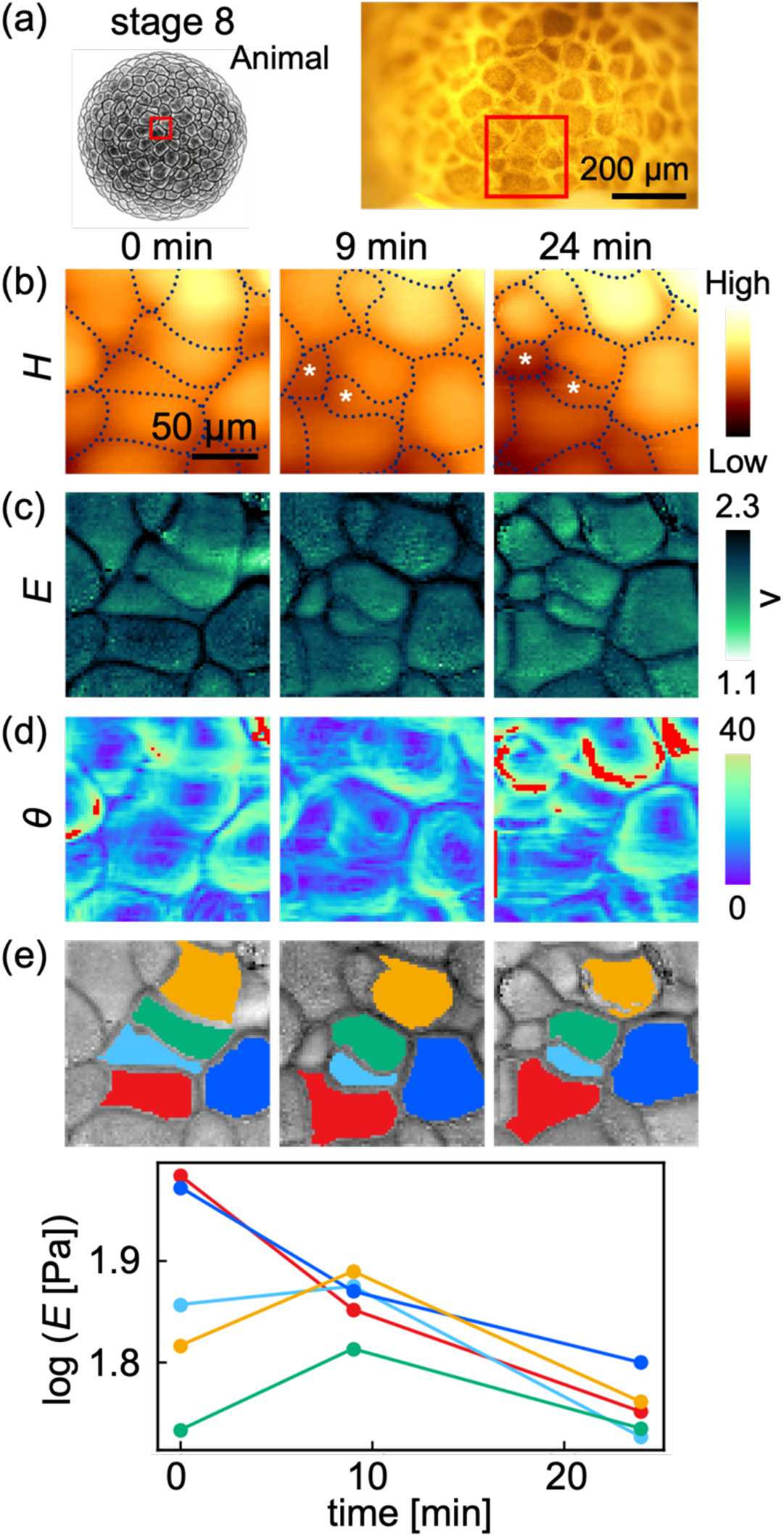
(a) Schematic morphology of *Xenopus laevis* embryo at stage 8 and an optical microscopic image, with AFM measured region indicated by an open square (red). Time-lapse mapping images of *H* (b), *E* (c), and *θ* (d) estimated from (b), where each mapping started at 0 min (left), 9 min (middle), and 24 min (right). Observed cell division is denoted by asterisks in (b). (e) A graph showing stiffness changes of individual cells over time. The colors in the graph correspond to those of the cells shown in AFM images in the upper figure.

Such characteristic mechanical behaviors of relatively large cells at the stage could be visualized, where more than 90% of the scanned regions exhibited surface tilt angles below 40° across fields exceeding 200 μm (Fig. 3d). The wide-range accessibility is essential for exploring dynamic mechanical behaviors over multicellular regions. For instance, we frequently observed dividing cell pairs undergoing asymmetric division (asterisks in Fig. 3b), where one daughter cell appeared substantially larger and topographically higher than the other (Fig. 3b). Such an asymmetric cell division is emblematic of *Xenopus laevis* embryos at stage 8, where blastomeres divide rapidly and often yield daughter cells of unequal size (*50*). Despite this morphological disparity, the two daughter cells showed no apparent difference in stiffness (Fig. 3c).

### 3.3 Stiffness in the animal hemisphere at the gastrula stage

At stage 10, *Xenopus laevis* embryos initiate gastrulation, which is characterized by coordinated movements of epithelial cells in the animal hemisphere directed toward the forming dorsal lip of the blastopore in the vegetal region (*30*). As gastrulation progresses toward stage 11, epiboly in the animal hemisphere becomes prominent. To capture the mechanical landscape during this critical phase, we performed AFM-based mapping of cells in the animal region at stage 10 (Fig. 4a). The results showed that the cells displayed polygonal morphologies (Fig. 4b), with different sizes and morphology (Fig. 4b, d). The cell stiffness in the apical cytoplasmic regions also varied substantially among individual cells (Fig. 4c, e), highlighting mechanical heterogeneity at the single-cell level. The cell stiffness was not correlated with the cell size (Fig. 4f). Consistent with earlier stages, we observed stiffening along cell-cell boundaries (Fig. 4c), which displayed a broad mechanical distribution, suggesting variations in cell-cell interactions.

**FIGURE 4.**
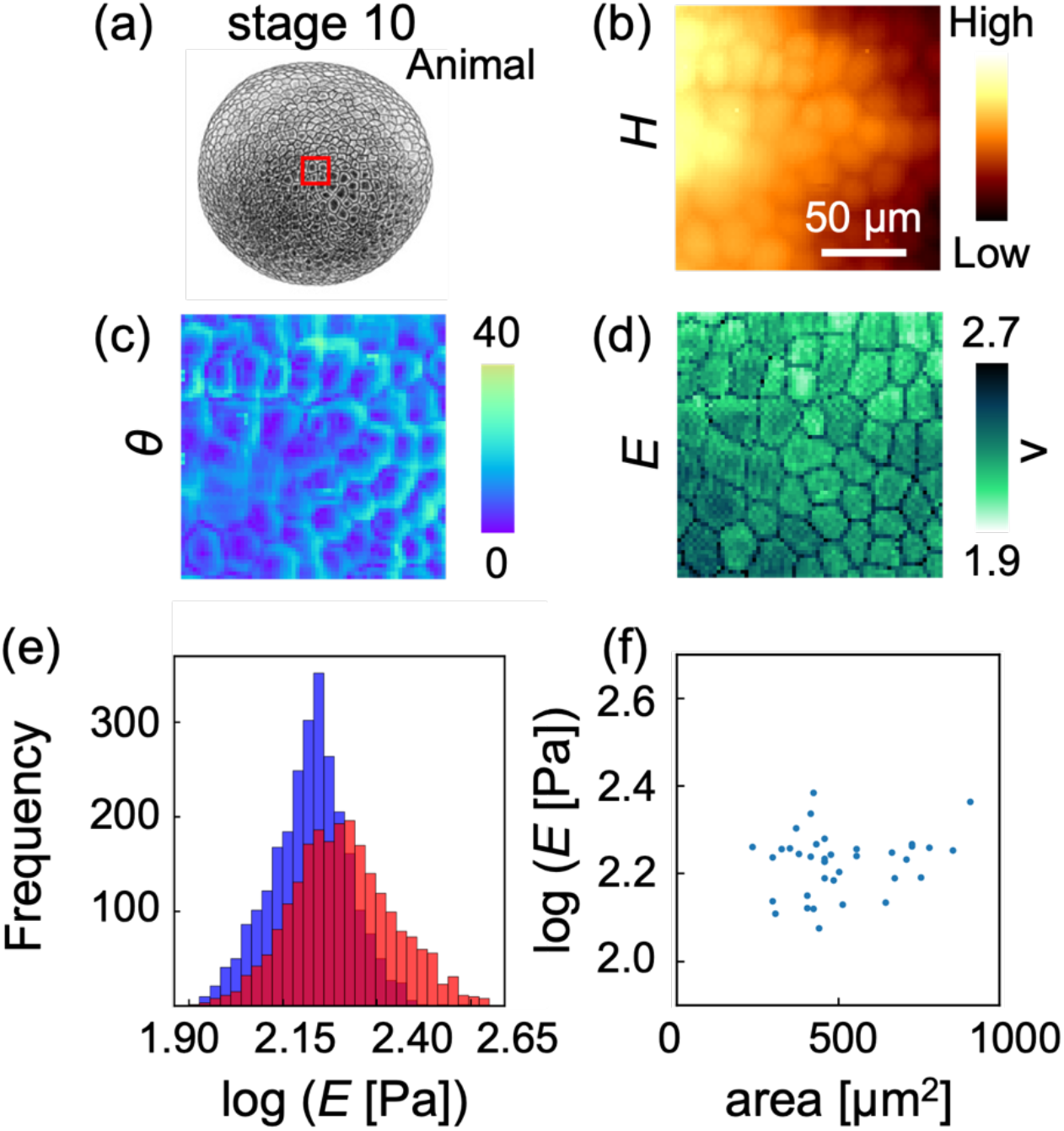
(a) Schematic morphology of *Xenopus laevis* embryo, viewed from the animal pole (Animal) and the vegetal pole (Vegetal) at stage 10. The AFM-measured region is indicated by an open square (red). AFM mapping images of *H* (b), *E* (c), and *θ* (d) estimated from (b). (e) Histograms of *E* values in the apical cytoplasmic region (blue) and the cell-cell boundary (red), shown in (c). (f) Plots of *E* values versus cell area (apical cell size) in individual cells, observed in (b) and (c), respectively. The coefficient of determination was 0.036.

### 3.4 Stiffness in the vegetal hemisphere at the gastrula stage

To investigate the mechanical properties of different germ layers, we conducted AFM mapping of *Xenopus laevis* embryos at stage 11 around the vegetal pole (Fig. 5a). The vegetal pole cells (yolk plug) displayed larger cell sizes (Fig. 5a, b) and less-rounded cell morphologies compared to cells in the animal hemisphere (Fig. 5d). Interestingly, AFM measurements revealed that cell-cell boundaries in the yolk plug were softer than the apical cytoplasmic regions (Fig. 5c, e), suggesting weak cell-cell interactions. Furthermore, in the apical cytoplasmic region, we observed discrete stiff granular structures without apparent changes in surface height (Fig. 5b), indicating localized mechanical heterogeneity within the cytoplasm (Fig. 5c). The average diameter of these granules was 6.9 μm (based on 158 granules measured in 19 cells from a single embryo). In contrast, cells in animal regions surrounding the yolk plug retained the canonical mechanical profiles in the animal hemisphere at earlier stages (Fig. 5b-e (III)).

**FIGURE 5.**
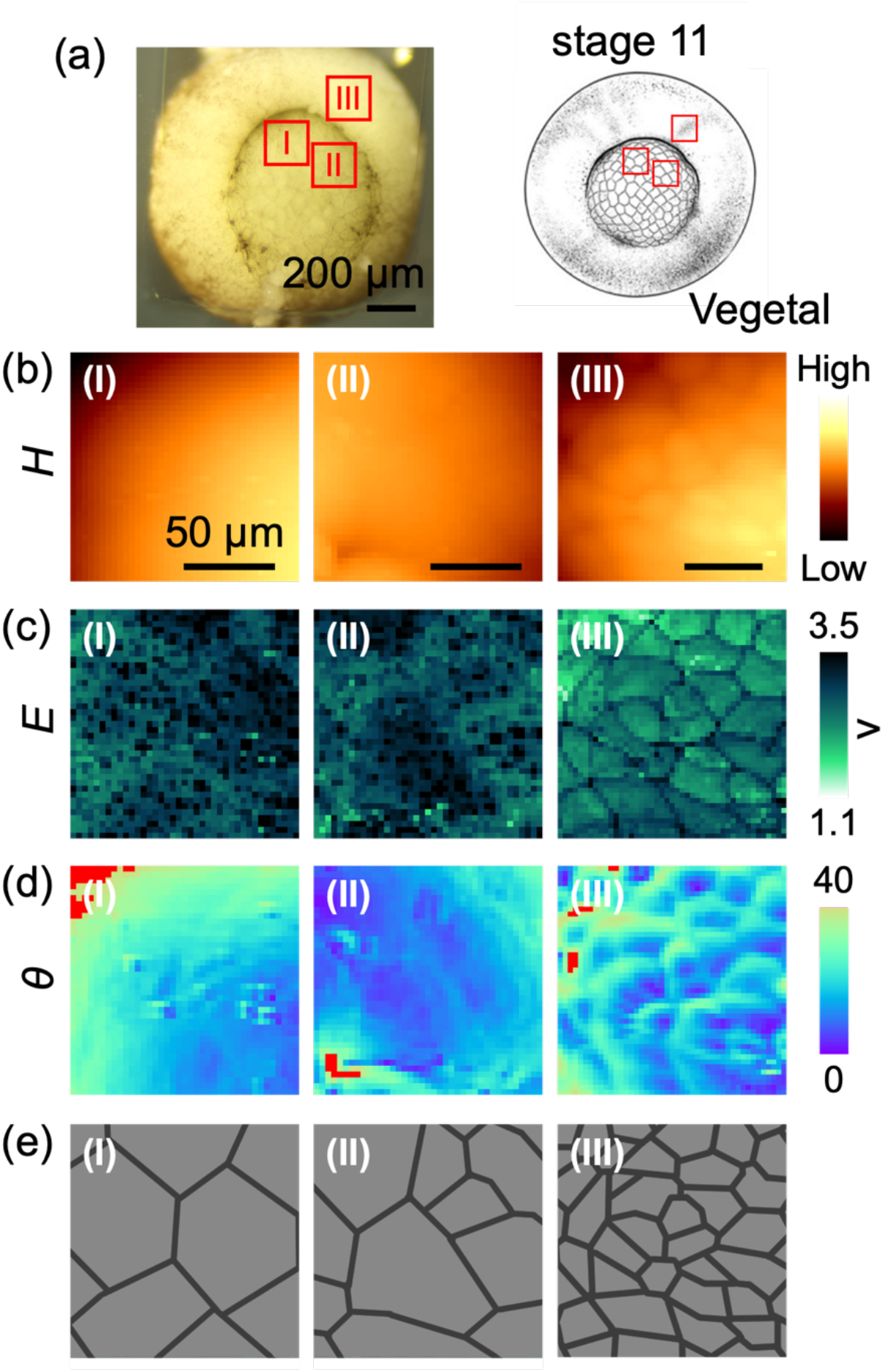
(a) An optical microscopic image and schematic image (inset) of *Xenopus laevis* embryo, viewed from the vegetal pole. AFM-measured regions are indicated by open squares: I and II are in the yolk plug, whereas III is in the animal region. AFM mapping images of *H* (b), *E* (c), and *θ* (d) estimated from (b). Schematic of cell shapes (e), estimated from (a), (b), and (d).

## 4. Discussion

### 4.1 Single-cell mechanics of *Xenopus laevis* embryo in the animal hemisphere

The present AFM data showed a variation in cytoplasmic stiffness during early *Xenopus laevis* embryogenesis (Figs. 2c, 3c, and 4c). We noticed that these stiffness values were similar to those reported in ascidian embryos during early cleavage stages (*23, 24*), underscoring a possible conserved mechanical origin across different species during early development. These results suggest that AFM can serve not only as a sensitive readout of biomechanical maturation within a single species but also as a comparative tool to dissect conserved physical principles underlying embryogenesis across species.

AFM mapping experiments revealed crucial mechanical signatures in the animal hemisphere: the stiffness along cell-cell boundaries was relatively higher than that in the cytoplasmic region, suggesting strong mechanical coupling between neighboring cells. Also, the cell stiffness exhibited marked cell-to-cell variability (Figs. 4c). The mechanical landscape of prospective epidermal cells in *Xenopus*, closely parallels that observed in epithelial monolayers in vitro (*43, 51*), suggesting that mechanical compartmentalization is an intrinsic feature of cohesive epithelial tissues, even in the complex three-dimensional environment of the *Xenopus laevis* embryo.

It has been shown that interfacial stiffness during cortical cell adhesion is governed by two principal factors: the elastic resistance of the cortical actin network and the dynamics of cytoskeletal remodeling (*52*). In the early low-contact phase of adhesion, cells establish initial contacts without substantial cytoskeletal reorganization. The cortical F-actin remains intact, maintaining elevated cortical tension and resulting in a mechanically rigid boundary. As adhesion matures via cadherin, which is a key regulator of epithelial cohesion, local cortical actin disassembles at the interface. This transition reduces cortical tension, thereby softening the boundary and permitting extensive membrane deformation and contact expansion. At stage 10, we detected a broad range of stiffness at the cell-cell boundaries (Fig. 4e), which may reflect heterogeneous cell-cell adhesion strength as well as localized variation in cytoskeletal organization.

An overtly asymmetric cell division, as shown in Fig. 3, exhibited a clear difference in apical cell height between daughter cells. This phenomenon may be interpreted from previous optical microscopy observations of *Xenopus laevis* embryos at stage 8 (*50*). In that study, cell division geometry is classified into three canonical configurations relative to the apical surface: parallel, perpendicular, or obliquely tilted (*50*). Based on this classification, we speculate that the asymmetric division observed by AFM (Fig. 3c) corresponds to an oblique configuration, in which the smaller daughter cell is displaced in the depth direction, thereby exhibiting a reduced cell height.

### 4.2 Single-cell mechanics of *Xenopus laevis* embryo in the vegetal hemisphere

The AFM mapping at stage 11 of *Xenopus laevis* embryo (Fig. 5) revealed distinct differences in mechanical properties between animal and vegetal hemispheres: cells in the yolk plug exhibited lower stiffness along cell-cell boundaries than in the cytoplasmic region, which was in contrast to cells in the vegetal hemisphere (Figs. 2*−*4). Previous studies revealed that E-cadherin is highly expressed in the embryonic ectoderm during gastrulation but notably absent in endodermal populations (*53*) (*54*). The distinct differences in stiffness pattern between cells in animal and vegetal regions observed by AFM (Fig. 5) may reflect the difference in cell adhesion protein expression.

Yolk platelets are distributed in the cytoplasmic regions (*55*). The size of platelets in cells changes depending on locations within the embryo and developmental stages (*56*). According to electron microscopy observations, the platelet size around the blastopore was around several micrometers in diameter (*56*). The size was comparable to that observed by AFM (Fig. 5c). The results suggest that AFM can be utilized for probing the microscale mechanical architecture of developing regulative embryos underneath the cell membrane in early vertebrate development.

## 5. Conclusions

Using force-indentation atomic force microscopy (AFM), we successfully quantified the spatial heterogeneity of mechanical properties in *Xenopus laevis* blastomeres in early embryogenesis at subcellular resolution. Removal of the vitelline membrane flattened the embryos and enabled direct measurement of embryonic cells with multi-probe cantilevers without perturbing normal development. AFM mapping revealed that cells in the animal hemisphere exhibited a mechanical profile sharply distinct from that in the vegetal hemisphere, indicating germ layer-specific mechanical signatures. The AFM measurements also detected subsurface features such as yolk platelets, underscoring its potential to probe intracellular structures relevant to developmental mechanics in *Xenopus laevis*. Overall, AFM offers a robust method for mechanical profiling of early *Xenopus laevis* embryos.

## Acknowledgments

We thank Dr. Kaori Kuribayashi-Shigetomi for assistance with editing the manuscript.

## Author contributions

T.M. and T.O. conceived and designed the experiments. M.Y., T.Y., T.K., Y.M., and T.M. prepared the samples. M.Y., T.K., Y.M., and T.O. performed AFM experiments and the data analysis. T.M. and T.O. supervised the experiments. M.Y., T.Y., T.M., and T.O. prepared the manuscript. All authors edited the manuscript prior to submission.

